# Spatial Transcriptomics of Glioblastoma Defines Spatially Anchored Transcriptional Activity Informing Therapeutic Vulnerability and Resistance

**DOI:** 10.1101/2025.10.17.683110

**Authors:** Visweswaran Ravikumar, Avery Maddox, Reva Kulkarni, Emre Kiziltug, Toshiro Hara, Arvind Rao, Wajd N Al-Holou

**Affiliations:** Department of Computational Medicine and Bioinformatics, University of Michigan, Ann Arbor, MI, 48109, USA; Department of Neurosurgery, University of Michigan, Ann Arbor, MI, 48109, USA; Biointerfaces Institute, University of Michigan, Ann Arbor, MI, 48109, USA; Department of Biostatistics, University of Michigan, Ann Arbor, MI, 48109, USA; Department of Radiation Oncology, University of Michigan, Ann Arbor, MI, 48109, USA

## Abstract

Intratumoral heterogeneity is thought to hinder targeted therapy in glioblastoma (GBM), but the extent of variation in known targets and tumor cell-intrinsic and extrinsic regulatory mechanisms within tissue remains unclear. In this work, we use Visium spatial transcriptomics to profile 44 tumor slides and 128,176 niche-annotated ST spots, including 14 new slides from 7 GBM specimens, along with spatial annotation from the Ivy Glioblastoma Atlas Project. First, we characterize the differential expression of *AVB3*, *EGFR*, *VEGF*, and *PDGFRA* pathway activities across space, elucidating the limited success of individual targeted therapies. Second, we identify transcription factor modules driving intratumoral heterogeneity and characterize a novel transcription factor module at the interface between core tumor and the invasive edge. Third, we identify cell type-specific receptor-ligand signaling enriched in specific spatial niches and relate these to specific molecular program alterations. Finally, we identify and characterize transcriptional pathways and their activity across spatial niches through transcription network analysis and demonstrate how these pathways can inform a combined therapeutic approach.

Together, through novel insights into the spatial patterning of transcription factor regulation, cellular interactions, and biological pathway activity, our work informs rational combination therapies targeting spatial niche specific vulnerabilities.

## Introduction

Glioblastoma (GBM) is the most common primary malignant tumor of the central nervous system (CNS), accounting for 48.6% of malignant CNS tumors, and one of the most lethal malignancies, with a median survival of only 15 months and a 5-year survival of only 7% [1,2]. Despite significant advances in the understanding of GBM and the identification of many novel therapeutic targets, the prognosis remains poor and the standard of care has not changed significantly in the last 15 years [2]. Trials of therapies targeting promising targets including *EGFR*, *PDGFRA*, *VEGF*, and *AVB3* have offered little to no improvement in GBM patient outcomes [3–5]. A major obstacle to advancing GBM therapy, indicated by its historical designation “Glioblastoma Multiforme”, is its extreme inter- and intra-tumoral heterogeneity [6]. Throughout and around the GBM tumor, distinct malignant and non-malignant cellular states, behaviors, and interactions exist in spatially defined microenvironmental niches that aid in the development of treatment resistance and tumor recurrence [6]. Elucidating this spatial heterogeneity is therefore critical to understand clinical failures, identify niche-specific targets, and develop combination therapy.

Significant work has been performed to explain the spatial patterning of GBM and provided important insights for improving treatment. In early work on this front, the Ivy Glioblastoma Atlas Project (Ivy GAP) integrated in-situ hybridization, laser microdissected RNA sequencing, and high-resolution imaging to define and map distinct GBM tumor regions including: leading edge (LE), microvascular proliferation (MVP), cellular tumor (CT) and pseudopalisading around necrosis (PAN) [7]. More recently, spatial transcriptomics (ST) analysis has revealed important GBM patterning in relation to these regions. The immune tumor microenvironment (TME) changes in relation to perivascular and perinecrotic regions and shapes resistance to immunotherapy [8]. Distinct glioma stem cell (GSC) signatures are enriched in relation to perinecrotic and peritumoral regions, marking region-specific increases in malignancy and infiltrative potential [9]. Further, distinct glioma cell states associate in neuron rich regions with unique vulnerabilities and invasion potential [10,11]. Additionally, a significant role for hypoxic stress gradients, which largely drive the patterning of Ivy GAP regions, has also been closely tied to the spatial patterning of GBM cell states, molecular behaviors, and treatment resistance [12,13]. Additional studies have helped spatially define the intercellular interactions in the GBM TME, and relate these interactions in their spatial niche to tumor progression and treatment resistance [14]. Despite these advancements, the extent of variation in potential and known therapeutic targets and their relation to tumor cell-intrinsic and extrinsic regulatory mechanisms within tissue remains unclear. With this work we sought to further elucidate the spatial patterning of these targets and their underlying regulatory mechanisms including cell-cell signaling and transcription factor programs.

Here, we generated a new Visium ST dataset of 14 slides from 7 GBM patients, and performed integrative analysis of our cohort with the Freiburg [15] and Broad [12] studies. Our integrated cohort includes 44 tumor slides and 128,176 niche-annotated ST spots after QC filters. Thus, our work presents findings from one of the largest GBM ST datasets analyzed to date, significantly enhancing confidence both where our study recapitulates prior findings as well as where we present novel results.

Differential expression analysis validates many previously described findings on the activation of distinct biological pathways and hallmark features across spatial niches and further demonstrates the limited spatial activity of targeted therapy-related pathways including *EGFR*, *PDGFRA*, *VEGF*, and *AVB3*. Analysis of tumor-specific expression profiles demonstrates continuous rewiring of tumor expression and tumor-specific transcription factor (TF) modules along spatial gradients including at the interface between spatial niches, evidencing localized vulnerabilities in the tumor cells. Analysis of cell-cell interactions identifies niche-specific ligand-receptor interactions including ephrin signaling at the leading edge (LE) between tumor cells and neurons and *AREG-EGFR* signaling between tumor and immune cells in core tumor (CT). We further identify and characterize spatially driven pathways in the form of co-expressed gene modules. At the hub of these pathways, we identify driver genes implicated in treatment resistance and vulnerable to targeted therapy. Further, we identify spatially defined programs enriched for specific drug targets, with evidence in the literature for activity against GBM.

Using a large cohort of Visium ST slides and cutting-edge analytical tools and techniques, we present a comprehensive overview of tumor biology across the diverse spatial niches in GBM. Our analysis reveals extensive tumor plasticity and variability of regulatory programs across spatial micro-environments, and elucidates why targeted therapies against single molecular markers are likely to be ineffective. Using our integrative analyses, we are able to highlight specific localized vulnerabilities and inform combination therapy development targeting these pathways.

## Results

### 1. Glioblastoma Anatomical Niches are Associated with Distinct Cancer Hallmark and Targeted Therapy Related Pathway Activity

We profiled 14 tumor slides from seven GBM patients on the Visium ST platform and pooled our data with additional Visium ST slides from two recent studies here termed the Broad (17 slides) and Freiburg (13 slides) cohorts [12,15] (Figure 1A). ST spots were annotated for spatial niches using the UCell algorithm [16] and Ivy GAP maker genes [7] (Figure 1B). After quality control, the combined dataset includes a total of 128,176 annotated spots with 65742, 8992, 33266 and 20176 spots from the CT, MVP, PAN and LE niches respectively. With sufficient representation of all niches (Figure 1C), we proceeded to characterize the changes in gene expression programs anchored by these niches.

**Figure 1:**
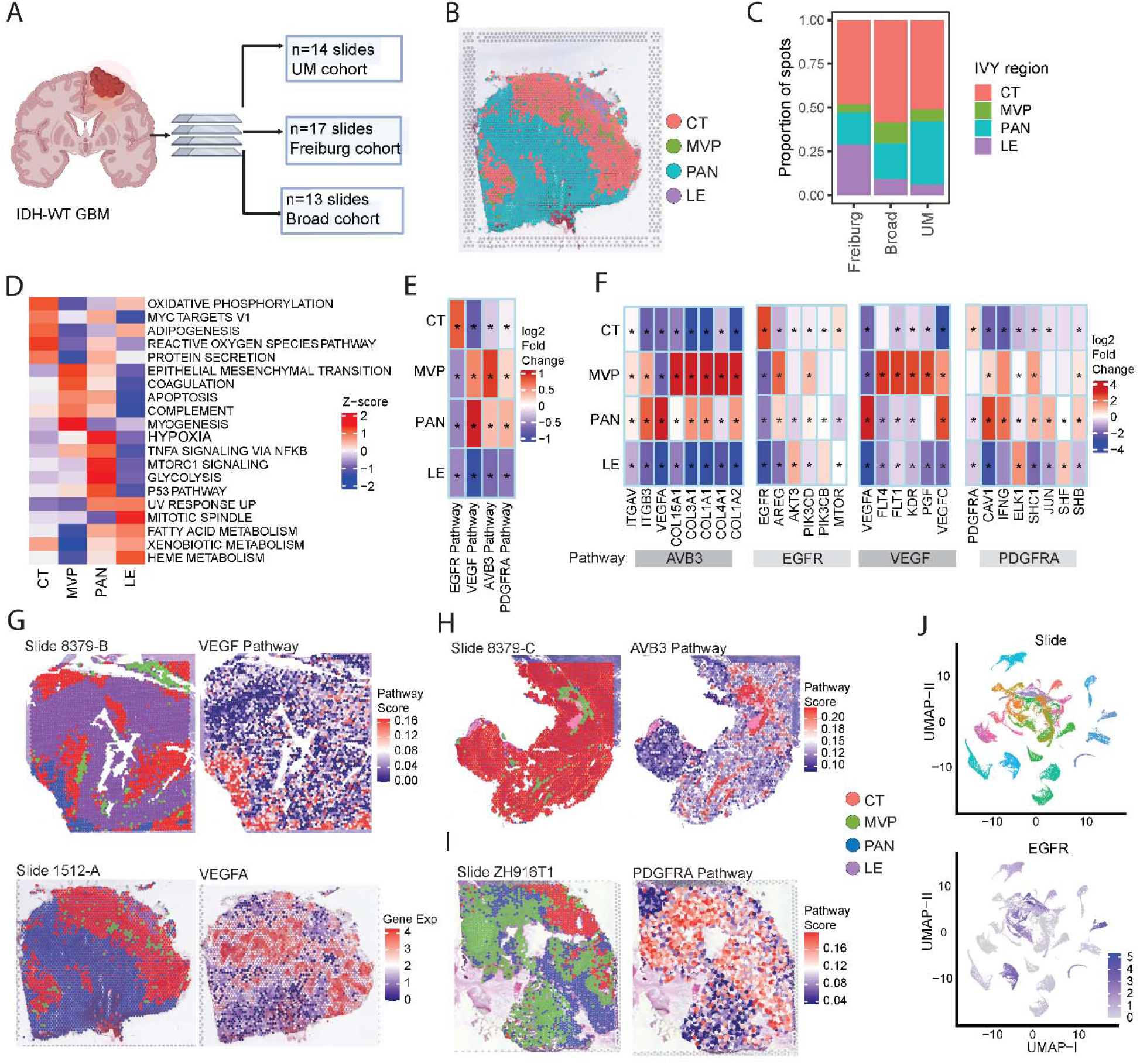
Pathway and drug target expression patterns of glioblastoma tumor microenvironments. **(A)** 14 GBM tumor specimen were collected in-house at the University of Michigan and combined with additional cohorts termed Broad (17 slides) and Freiburg (13 slides) [12,15]. **(B)** Ivy GAP marker-based annotation of anatomical niches. **(C)** Relative niche composition across study cohorts. **(D)** Top-5 differential Hallmark pathways per niche. **(E)** Differential expression analysis of *AVB3*, *EGFR*, *VEGF*, and *PDGFRA* pathway activity and **(F)** top differentially expressed genes for each pathway in addition to titular pathway genes. For **E** and **F**, * where FDR < 0.05. **(G-I)** Representative spatial visualization of **(G)** *VEGF,* **(H)** *AVB3*, and **(I)** *PDGFRA* pathway activity. **(J)** UMAP of combined cohort showing inter-patient heterogenous expression of *EGFR*.

Differential expression analysis across tumor niches identified 2,642 significant genes, with the LE exhibiting the most distinct transcriptional profile. Gene ontology (GO) term [17] enrichment analysis of these genes revealed biologically coherent patterns (Supplement Figure 1C). Vasculature development and angiogenesis pathways were enriched in MVP, while hypoxia-response signatures predominated in PAN. Across the tumor bulk, we observed significant enrichment for extracellular matrix (ECM) activity. CT regions showed upregulation of neuronal activity, glial cell differentiation, and Notch signaling. Finally, LE was uniquely enriched for neuronal signaling and synaptic activity, indicating the integration of tumor cells with surrounding neural networks [18].

Comparison of enrichment scores for cancer hallmark signatures demonstrated well supported niche-specific roles (Figure 1D). CT was enriched for protein secretion, a known hallmark of the GBM tumor core where non-canonical pathway secretion of ECM proteins and soluble proteins like *CD109* promotes tumor cell motility, invasiveness, and disease progression [19–21]. The MVP and PAN regions shared enrichment for complement signaling and coagulation pathways, which are associated with functions including vascular remodeling, modulation of immune activity and supporting tumor cell survival [22–24]. Epithelial mesenchymal transition (EMT) activity was enriched in and around the MVP niche (Supplement Figure 1D).

The LE was enriched for mitotic spindle activity, a hallmark closely linked to both proliferation and invasiveness. LE and PAN regions were enriched high UV response activity, suggesting a robust DNA damage response [25]. While the PAN niche has a high capacity for sensing DNA damage, downregulation of DNA-damage repair mediators result in accumulated chromosomal alterations which contribute to subclonal evolution of the tumor [15,26]. In contrast, the LE has robust DNA repair activity making infiltrative cells resistant to radiation and chemotherapy, driving tumor recurrence [27]. LE was also enriched for adipogenesis, fatty acid oxidation and heme metabolism processes which have been shown to facilitate the adaptability of cancer cells to their environment, supporting their survival and invasion [28–30]. Altogether, these findings robustly confirm the spatially defined activity of several previously described niche-specific GBM programs.

This spatially patterned niche-specific transcription sheds light on the limited advancements realized through targeted therapies for GBM. Relapse remains a practical inevitability and there has been no significant improvement in survival since the Stupp protocol [31] of surgical resection, chemoradiation and adjuvant chemotherapy with temozolomide (TMZ) became standard of care in 2005 [3]. As GBM cells are highly plastic and versatile owing to their clonal evolution from GSCs with significant differentiation and dedifferentiation capabilities [32,33], it unlikely that any single therapeutic modality would be effective in treating the entire heterogeneous population [3]. This is well demonstrated by the ineffective outcome in trials of therapies developed against promising targets including *EGFR*, *PDGFRA*, *VEGF*, and *AVB3* [3–5].

Differential expression analysis of the activity in these target pathways demonstrates their localization to specific niches, and emphasizes their limited activity in the LE (Figure 1E). This variability is further emphasized when inspecting individual pathway genes, demonstrating additional spatially patterned alterations in pathway component behavior (Figure 1F).

Representative slides visualize the strong localization patterns of these pathways including the *VEGF*-pathway to PAN, *AVB3*-pathway to MVP, and *PDGFRA*-pathway to MVP and PAN (Figure 1G-I). In addition, UMAP of *EGFR*, which is reported amplified in 57.4% of primary GBMs [3], across the combined cohort recapitulates its interpatient heterogenous expression pattern in GBM (Figure 1J). Together, these findings help to link the heterogeneous expression patterns of these target pathways and their constituent genes with the ineffective outcomes observed in clinical trials.

### 2. Transcription Factor Activity Along Spatial Niche Gradients Drives Continuous Molecular Reprogramming in Tumor Cells

We next investigated the TF orchestration of tumor cell-specific transcriptional reprogramming across the tumor niche continuum. We employed *BayesPrism* [34] to extract tumor cell-specific expression data for each spot (Figure 2A). The resulting tumor-expression matrix was normalized and used to investigate spatial variation in gene expression across anatomical niches.

**Figure 2:**
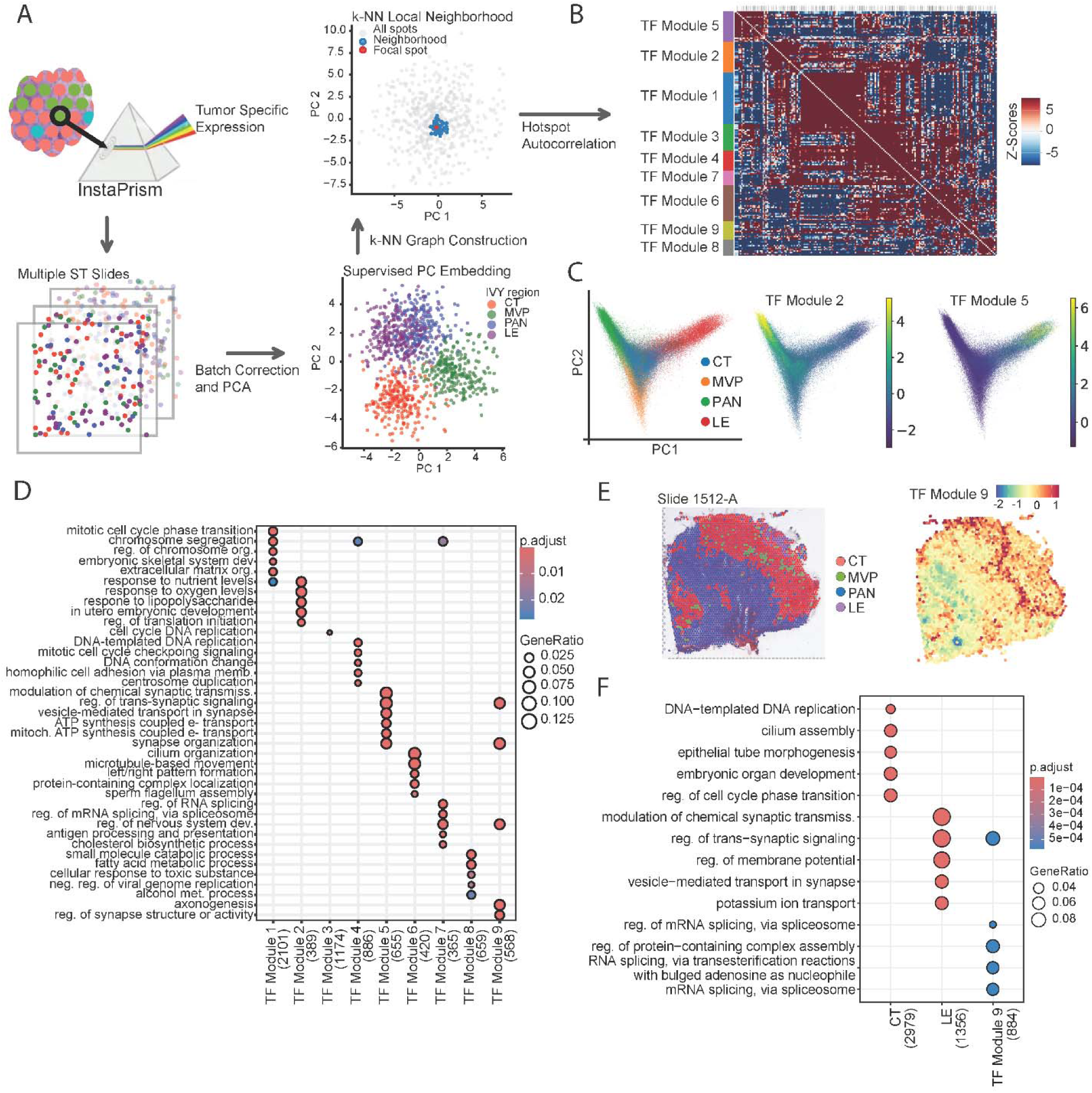
Spatially driven adaptation of tumor cells in GBM. **(A)** Schematic of hotspot network construction. Tumor specific expression was extracted from ST data, corrected for batch effect, and underwent PCA. Autocorrelation was calculated on neighborhoods of spots identified through k-NN graph construction in PC embedding. **(B)** This Hotspot analysis identified 9 distinctly autocorrelated TF modules with spatially localized activity across all slides. **(C)** Localized activity scores of representative TF modules in PC space. **(D)** Enrichment analysis of differential genes in spots enriched for specific TF module activity. **(E)** Spatial score of TF Module 9 on representative slide shows localization to CT-LE interface. **(F)** Comparative analysis of TF Module 9 enriched spots with CT and LE identifies biological processes specifically active in the CT-LE interface.

Differential expression analysis of tumor-expression data identified genes and TFs exhibiting significant changes between spatial niches. Recognizing that tumor spots exist along a spatial continuum, we generated a supervised PC embedding using batch-corrected expression for the Ivy GAP marker genes in each spot. This low-dimensional embedding succinctly recapitulates the major axes of transcriptional variation between tumor niches (Supplement Figure 2A). We leveraged the relative enrichment scores derived from this embedding to select the top 3,000 scoring spots for each niche.

Differential expression analysis identified a total of 3,700 genes, underscoring the extensive transcriptional rewiring that occurs as tumor cells adapt to local spatial environments. Functional enrichment analysis of these genes (Supplement Figure 2B) revealed region-specific biological programs: CT was enriched for DNA replication, cell cycle regulation, differentiation, and cell motility; MVP region showed upregulation of ECM reorganization processes; PAN was characterized by enrichment of hypoxia response, apoptotic signaling, and wound healing; and LE was characterized by synapse organization and signaling. These findings are consistent with prior reports that the mesenchymal (MES) state in hypoxic GBM regions recapitulates an injury-response inflammatory signature [35]. In the LE, invasive tumor cells exhibit upregulation of genes indicating a functional integration with the surrounding neuronal microenvironment [36]. In parallel, we identified 343 TFs with significantly altered expression across spatial niches, with 246 showing highest expression in CT, highlighting the regulatory complexity within this compartment (Supplement Figure 2C).

To investigate TFs underlying tumor cell-specific reprogramming, we adopted the *Hotspot* [37] algorithm, a graph-based method for detecting genes with localized expression patterns in single-cell and spatial data. We constructed a k-nn neighborhood graph using our unified PC embedding, and systematically assessed the expression of human TFs [38] in this space (Figure 2A). *Hotspot* identified 184 TFs with significant local autocorrelations (FDR < 0.05), which were split into nine modules of covarying TFs (Figure 2B). TF modules showed niche-specific activation: TF Module 2 was distinctly enriched in PAN domains, while TF Module 5 was active in the leading edge (Figure 2C). Spatial activity maps (Supplement Figure 3) revealed that most TF modules (TF Module 1, 3, 4, and 7) are concentrated around the perivascular niche, indicating substantial and distinct regulatory processes in this region. TF Modules 8 and 9 were predominantly active along the CT-to-hypoxic and CT-to-LE transition zones, respectively. Correlation analysis of module activity across all spots (Supplement Figure 2D) recapitulated the spatial segregation of regulatory programs. TF Module 8 and 9 were weakly correlated with the CT TF modules and anticorrelated with each other, highlighting their distinct roles along different spatial interfaces. These results further enforce how specific TF modules orchestrate localized gene expression programs as tumor cells adapt to their microenvironment [39,40].

To assess the regulatory significance of these TF modules, we selected the top 1,000 highest-scoring spots for each TF module and compared their gene expression profiles. Although many TF modules showed peak activity in the peri-vascular region, their enrichment was spatially distinct, resulting in minimal overlap among the top spots across modules. This spatial segregation suggests a fine-grained organization of biological processes, particularly within perivascular regions, consistent with recent studies demonstrating niche-specific regulatory programs and the compartmentalization of tumor-cell phenotypes [8,10,15].

GO enrichment analysis (Figure 2D) revealed that each TF module is associated with a distinct set of biological processes. TF Module 1, 3, and 4 are primarily linked to cell division and DNA replication, reflecting proliferative programs spatially confined within the tumor core and perivascular regions [10]. TF Module 6 is specifically enriched for cytoskeletal organization and cell motility, aligning with the known roles of perivascular niches in supporting tumor cell migration and invasion [41,42]. TF Module 7 shows distinct enrichment for RNA splicing processes. Alternative splicing is emerging as a critical disease mechanism in GBM, generating transcript diversity that contributes to tumor heterogeneity and impacts neural developmental hierarchies [43,44]. Co-localization of activity of these diverse TF modules around vascular structures underscores the multifaceted nature of perivascular tumor adaptation.

PAN-specific TF Module 2 is enriched for pathways related to adaptation to hypoxia and nutrient deprivation, as well as developmental processes indicative of GSC maintenance. These findings are consistent with the immunosuppressive and stem-like character of the perinecrotic niche [13,45]. TF Module 5, active at the LE, is enriched for neuronal signal transduction, synaptic signaling, and mitochondrial ATP synthesis, suggesting that tumor cells at the invasive margin are both highly metabolically active and potentially integrating into neural circuits.

TF Module 8, which is active at the CT–PAN interface (Supplement Figure 3), is associated with diverse metabolic processes, reflecting metabolic reprogramming as tumor cells transition toward hypoxic environments. TF Module 9, localized to the CT–LE interface (Figure 2E), shows enrichment for neurodevelopmental processes, synapse organization, and axonogenesis, indicating that tumor cells in this region may be acquiring neuronal phenotypes as they prepare to invade the brain parenchyma.

TF Module 9 emerged as a novel regulatory hub for adaptations along the CT-LE gradient. To delineate biological mechanisms specific to this interface, we performed differential expression analysis specifically comparing gene expression along this gradient. We selected the top 1000 scoring spots for the CT and LE Ivy GAP markers, and 1000 spots with strongest TF Module 9 activity. This revealed 940 genes significantly upregulated in TF Module 9 spots, with pronounced enrichment for mRNA splicing/processing and neurodevelopmental pathways (Figure 2F). The alternative splicing machinery is overrepresented, aligning with the demonstrated role of this molecular mechanism in inducing neural stem-like states in GSCs and enhanced adaptability to microenvironmental challenges [44,46]. These findings implicate post-transcriptional regulation as a critical driver of invasive phenotypes, suggesting targeting spliceosome components as a potential strategy to contain tumor spread.

The relative activity of individual TF modules varies substantially between slides (Supplement Figure 2E), reflecting the underlying heterogeneity in niche composition observed between slides (Supplement Figure 1B). This variability is consistent with the highly localized and niche-specific activation of TF modules, many of which are confined to distinct spatial interfaces that may only be present in a subset of samples. This highlights the necessity of profiling a large and diverse cohort to capture the full spectrum of spatial regulatory heterogeneity in GBM. Further, these TF programs underly the spatial adaptations and heterogeneity of established pathogenic GBM cell programs, making them promising therapeutic targets in combination with downstream pathways.

### 3. Cell type composition and ligand-receptor signaling are unique across spatial niches

To resolve TME cellular architecture, we performed reference-based deconvolution on every ST spot using a harmonized GBM SC atlas [47] and *InstaPrism* [48] (Figure 3A). The inferred cell type abundance matrix was modeled as compositional data and used to construct a sparse cell-interaction graph (Supplement Figure 4A). Consistent with the concept of a layered organization of the GBM TME [12], we see that cell types have limited positive interactions with other TME components, alongside numerous negative interactions indicating localized distribution in distinct spatial regions. We further visualize niche specific TME composition patterns (Figure 3B). As with gene expression programs, we see a clear transition between tumor niches. We also note strong localization of different Neftel tumor states to specific niches (Figure 3C), with the NPC-like cells localized to the LE; AC- and OPC-like cells most prevalent in the perivascular niches; and MES-like tumors most abundant in the hypoxic areas. Our observation is consistent with known localization patterns of these tumor states to specific TME niches, and indicates the power of spatial context in the TME to shape cellular expression programs [49].

**Figure 3:**
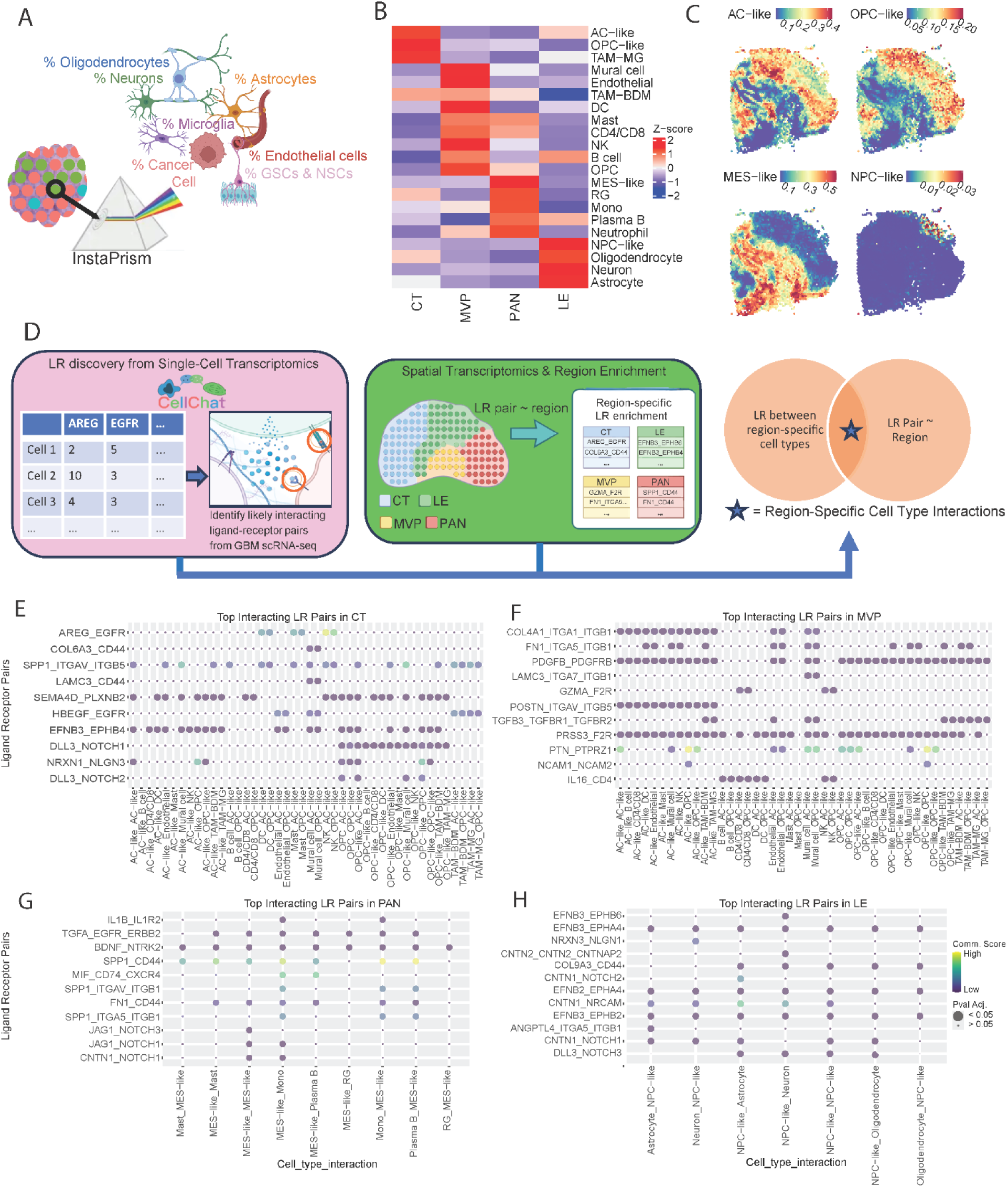
Cell type composition and ligand-receptor analysis reveal distinct signaling and interaction patterns across spatial niches. **(A)** BayesPrism resolves spot-level expression data into cell type composition. **(B)** Scaled abundance of cell types in tumor niches. **(C)** Localization of Neftel tumor states to distinct spatial niches in representative slide. **(D)** IVY Gap niche specific LR interactions are obtained using both SC and ST GBM data. **(E)** CellChat communication scores for top LR pairs. LR pairs shown whose spatial co-expression is associated with each niche. Note, as scores are dependent on expression values, scoring is relative across cell type combinations and LR pairs.

We next sought to characterize continuous transitions in expression programs along spatial niche gradients using our low-dimensional embedding, which succinctly recapitulates the major axes of transcriptional variation between tumor niches (Supplement Figure 2A).

Enrichment scores for the Neftel state [50] marker genes in the PC embedding demonstrates fine-scale distinctions in relative localization patterns of the MES1/MES2 and NPC1/NPC2 tumor cell states (Supplement Figure 4B). The MES2 signature was specifically enriched in extreme PAN regions, consistent with its established association with hypoxia-responsive programs in GBM [39,50]. Within the NPC-like compartment, the NPC2 program localized to the tumor leading edge, identifying highly proliferative, progenitor-like populations in the invasive margin, while the NPC1 signature was prominent in the cellular tumor core, consistent with a more differentiated, less proliferative state [51].

We next aimed to decipher niche-specific cell signaling through integration of ST and SC ligand-receptor (LR) analyses, and filtered results for niche-represented cell types (Figure 3D). *CellChat* [52] scores for select LR pairs in each niche (CT, MVP, PAN, LE), ordered first by unique spatial association with the region and second by modeling coefficient, demonstrate the unique signaling behavior between niches (Figure 3E-H).

We start by investigating the CT niche. Top interactions include *AREG*-*EGFR*, and ECM-*CD44* signaling with *COL6A3* and *LAMC3* ligands. *EGFR* is a known oncogene across cancers, where overexpression of this protein promotes uncontrolled cell proliferation through the activation of downstream pathways such as *PI3K*, *MAPK*, and *STAT3* [53–55]. A recent glioma study proposed the role of tumor-associated monocytes (TAMo’s) in driving mesenchymal transformation of tumor cells through the *AREG*-*EGFR* signaling axis [56]. In the CT niche, our analysis shows that the *AREG*-*EGFR* interaction is most likely mediated by DC, Mast, or NK cells, targeting OPC-like or AC-like tumor states (Figure 3E).

Both *LAMC3* and *COL6A3* are ECM components and their interaction with *CD44* in normal tissue is primarily to maintain tissue structure [57]. In the context of the *COL6A3-CD44* pair, *COL6* protein has been found significantly upregulated in GBM and strongly correlates with a stem-like phenotype and chemotherapy resistance [58,59]. Similarly, another study indicates the role of laminin alpha-2 in GSC maintenance and growth [60]. Our findings indicate both *LAMC3*- and *COL6A3*-*CD44* pairs most likely interact between Mural Cells with AC and OPC-like tumor cells (Figure 3E).

In the MVP niche, interactions of interest include integrin signaling (*COL4A1*-*ITGA1*-*ITGB1*), *GZMA*-*F2R*, and *PTN*-*PRPZ1*. Integrin signaling is consistently activated in the MVP region by different ECM-ligands including collagen, *FN1*, laminin, and *POSTN* (Figure 3F). In GBM, many studies identify the critical pro-angiogenic role of integrins, discovering high expression of these receptors on endothelial cells [61,62]. Similarly, our highest *CellChat* communication scores for integrin signaling are between mural-mural or mural-endothelial cells (Figure 3F), likely providing a scaffold for endothelial cell migration when creating new vessels. In addition, we note that among all tumor-specific interactions, *COL4A1*-*ITGA1*-*ITGB1*, along with *FN1* and *LAMC3* interactions, are most likely between mural and OPC-like cells (Figure 3F). Supporting this interaction, an integrin-driven increase in tumor invasive properties has been identified in GBM [63]. Thus, our findings reinforce angiogenesis driven by integrin signaling on endothelial and mural cells in MVP.

*GZMA* is a protein found in NK cells and cytotoxic T cells, playing a role in target cell death [64]. A recent study proposed *GZMA-F2R* interaction played an anti-tumorigenic role in hepatocellular carcinoma (HCC) where *GZMA* secreted by T-cells interacts with *F2R* expressed by tumor cells to induce T-cell mediated tumor suppression via activation of the *JAK2*/*STAT1* pathway [65]. In our analysis, among tumor state interactions in MVP, this pair most likely interacts between CD4/8 or NK cells as sender, and AC-like or OPC-like tumor states as receiver (Figure 3F).

Several studies identify *PTN*-*PTPRZ1* interaction between TAM’s and GSCs by showing that silencing *PTN* expression in TAMs mitigates their pro-tumorigenic activity [66,67]. These studies reveal *PTN* binding to *PTPRZ1* allows GSCs to self-renew and maintain stem-like properties. However, our findings indicate this LR pair most likely interacts between AC- and OPC-like cells (Figure 3F), suggesting an alternative signaling route in the MVP region. In recent works, *PTPRZ1* was identified as a marker gene for an OPC-like enriched transcriptomic cluster, specifically emphasizing the expression of *PTPRZ1* along orG differentiation to OPC-like cells in GBM [68,69]. While there is no direct literature on *PTN* presence in AC-like tumor state, studies indicate this protein is expressed by both GBM tumor cells and *TAMs* [70].

Analysis of the PAN niche identifies specific *CD44* signaling (*SPP1*-*CD44*, *FN1*-*CD44*) and *MIF*-*CD74*-*CXCR4* interactions. *MIF* is a cytokine that can bind to both *CD74* and *CXCR4* as a complex, typically triggering leukocyte modulation and recruitment [71]. Similarly, our analysis indicates this pair most likely interacts between MES-like cells and monocytes (Figure 3G). Prior work reveals that across cancers, the *MIF*-*CD74* axis acts through immune suppression, modulating classical/alternative polarization of monocytes [72–74]. However, the role of *MIF* in GBM is complex, presenting both anti and pro-tumorigenic effects depending on the context [75]. The over-expression of *MIF* in response to hypoxic stressors has been shown in several studies [76,77] and hypoxia-induced vascular mimicry (VM) has been proposed through the *CXCR4*-*AKT*-*EMT* pathway driven by *MIF* [77].

The interactions of both *FN1* and *SPP1* with *CD44*, identified in PAN, are typically involved in wound healing processes, in contrast to the ECM component interactions with *CD44* identified in CT, whose primary role is tissue integrity [78]. Macrophage signaling via the *SPP1*- *CD44* axis across cancers drives T-cell exhaustion and contributes to immunosuppression [79–81]. However, our findings indicate these *CD44* interactions in the PAN region are most likely between monocytes and MES-like tumor cells, as T-cells are not enriched in this region (Figure 3G). Supporting an alternative mechanism of *SPP1*-*CD44* interaction in high-grade glioma, prior work has proposed that this interaction plays a critical role in macrophage-mediated induction of MES-like tumor state [82]. Thus, we hypothesize a niche-specific mechanism of these *CD44* pairs, particularly *SPP1*-*CD44*, in driving MES-like tumor state induction in the PAN region through macrophage-tumor signaling.

Finally, top interactions in the LE niche include ephrin signaling and *CNTN2*-*CNTN2*-*CNTNAP2*, and *CNTN1*-*NRCAM*, which focus different aspects of neuronal development including axon guidance and synaptic growth and maintenance [83,84]. In GBM, ephrin interactions have a complex role, with some promoting tumor-suppressive behavior such as *EphA1* and *EphA5*, while many promote aggressive invasion particularly along white matter tracts such as *EphA2*, *EphA3*, and *EphB3* [85,86]. Our findings in the LE show the strongest ephrin pair interaction, *EFNB3*-*EPHB6*, between NPC-like and neuron cells (Figure 3H). *CNTN2*-*CNTN2*-*CNTNAP2* supports neuronal development by forming a synaptic bridge, particularly at the nodes of Ranvier [87]. Consistent with this function, our findings emphasize this interaction between NPC-like cells and neurons (Figure 3H).

Taken together, these results advance the understanding of cell signaling in GBM by coherently placing established interactions into specific spatial contexts. Further, these results highlight the niche-specific scope of key interactions, a major limitation to any single targeted therapy and key consideration for a combined therapeutic approach.

### 4. Gene Co-Expression Network Analysis Characterizes Niche-Specific Pathway Activity, Drivers of Spatial Program Patterning, and Spatially Defined Therapeutic Targets in GBM

We further sought to identify and characterize distinct spatially driven molecular programs directly from our ST data and leverage them towards identifying niche and gene-specific therapeutic targets. Towards this end, we employed gene co-expression network (GCN) analysis on our ST dataset using *hdWGCNA* [88]. In brief, *hdWGCNA* aggregates proximal clusters of ST spots within each niche into metaspots, computes the co-expression between genes across metaspots, and clusters genes by their co-expression similarity into distinct modules (Figure 4A). Our GCN analysis identified 19 modules, labeled in order of decreasing size (M1-M19). Each module putatively represents a distinct program or closely associated set of programs.

**Figure 4:**
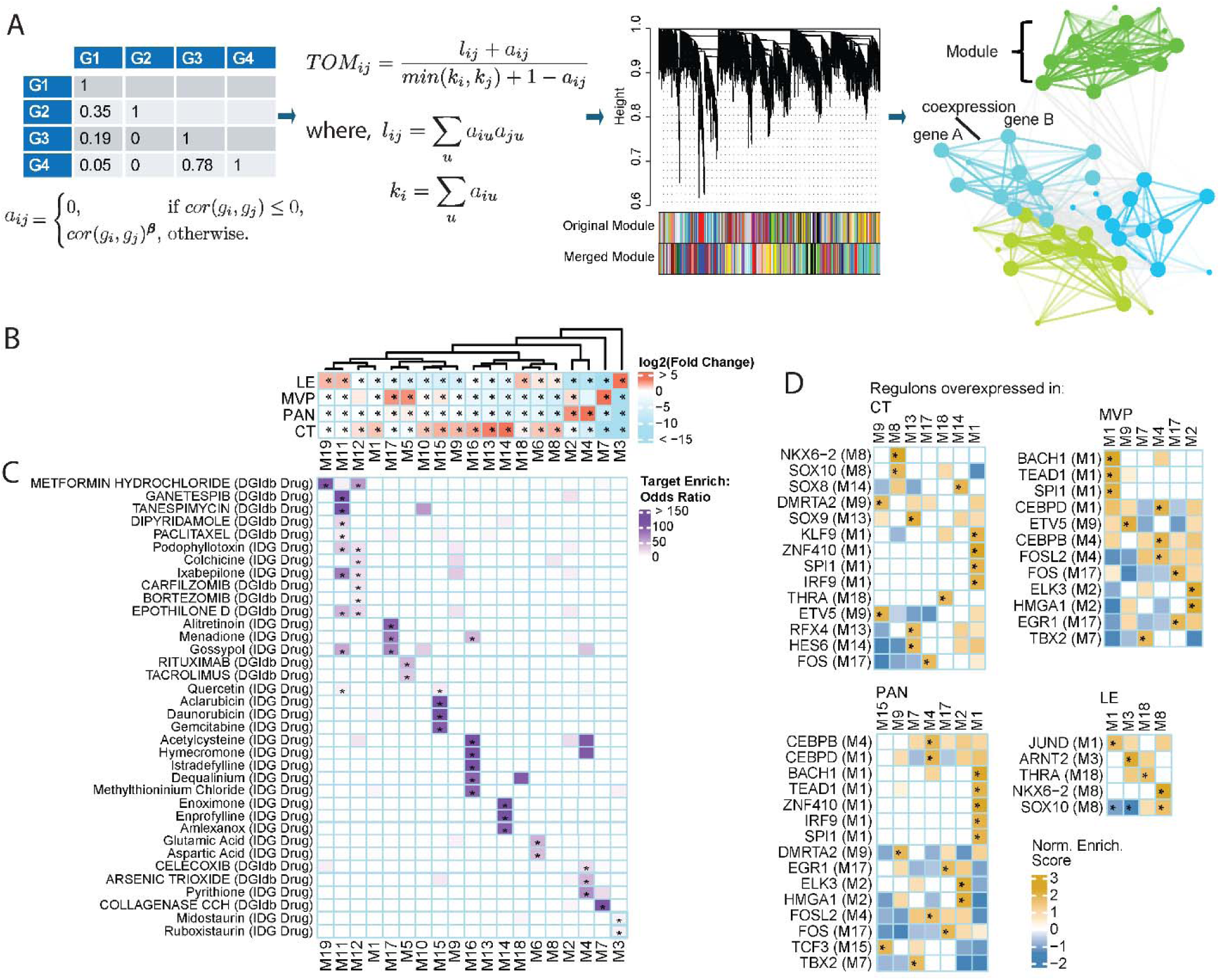
Co-expression network analysis of GBM ST data characterizes spatial niche-driven pathway dysregulation, core drivers, and candidate drug vulnerabilities. **(A)** Summary of network construction including similarity adjacency matrix calculation from metaspot gene expression data, topological overlap matrix (TOM) calculation, hierarchical clustering of genes on TOM dissimilarity, and resultant network with genes as nodes and coexpression similarity as edges. **(B)** One-vs-all testing of module eigengenes in each niche relative to rest of tumor. **(C)** Drug target enrichment results in each module. **(D)** Module enrichment for niche-associated regulon transcription factor targets. * = FDR < 0.05.

Module eigengenes (MEs), the first principal component of a module’s gene expression profile, provide a single continuous value capturing the activity level of a module. MEs can then be modeled against IVY Gap niches to identify spatially driven programs (Figure 4B, Supplement Figure 7). Modules can be further characterized through enrichment analysis for pathway terms (Supplement Figure 5A) and inspection of their leading hub genes, the genes most connected within their module and candidate drivers of module behavior (Supplement Figure 5B). Spatially defined therapeutic targets can be identified through module enrichment for drug targets (Figure 4C). Further, we constructed a transcription factor regulatory network (TRN) from our ST data using *hdWGCNA*. TRN regulons represent individual TFs and their strongest negatively and positively regulated targets. Through enrichment analysis of GCN modules for regulon TF targets (Figure 4D), we nominate key TFs driving the spatial programming of pathway activity in GBM. Finally, we modeled ME expression against LR coexpression to identify signaling interactions driving module behavior (Supplement Figure 6A).

M14, M13, M16, and M15 were the most overexpressed modules in CT (Figure 4B). M14 is enriched for NOTCH signaling pathways (Supplement Figure 5A), reinforcing earlier findings (Supplement Figure 1C). Leading M14 hub genes *HES6*, *MARCKSL1*, *OLIG1*, and *BCAN* (Supplement Figure 5B) closely relate it to the NPC1 tumor state [50], recapitulating prior findings of NPC1 tumor state enrichment in CT (Supplement Figure 4B). Further, M14 is associated with *COL6A3-CD44* and *LAMC3-CD44* signaling, supporting a role for these interactions in GSC maintenance in CT through AC- and OPC-like tumor cells (Supplement Figure 6). Interestingly, PDE inhibition has shown promise for GBM therapy [89,90] and M14 is enriched for targets of 3 closely related PDE inhibitors: Enoximone, Milrinone, and Amrinone (Figure 4C). Together these highlight a specific NPC1-like niche that may be targeted through PDE inhibition.

M13 is enriched for glial cell differentiation, negative regulation of neuron differentiation, and transport across the blood-brain barrier (Supplement Figure 5A). Leading hub genes *MLC1*, *PON2*, *GATM*, *EDNRB*, *TTYH1* and *ATP1B2* (Supplement Figure 5B) are markers of the Neftel AC state [50], reinforcing earlier findings of the AC tumor state enrichment in CT (Supplement Figure 4B). Hub gene *PON2* has been identified as a target of valproic acid (VPA), playing a central role in reactive oxygen species (ROS) regulation that is acted upon by VPA leading to increased ROS production and GBM growth inhibition [91]. M13 is enriched for positively regulated targets of *SOX9*, *RFX4*, and *HES6* (Figure 4D) which are all overexpressed regulons in CT (Supplement Figure 8). These TFs have been identified as core regulators of GBM programs [92–94]. Of note, *SOX9* is a driver of radiation and temozolomide (TMZ) therapy resistance and is being researched as a target in a novel *SOX9*-responsive suicide gene therapy for GBM [94].

M16 is enriched for the regulation of amyloid-beta formation, amide metabolic process, and cellular component biogenesis (Supplement Figure 5A). M16 hub genes *CST3*, *S100B*, *GFAP*, *AQP4*, and *CLU* (Supplement Figure 5B) are all markers of the Neftel AC tumor state [50], recapitulating prior findings of AC-like cell enrichment in CT (Supplement Figure 4B).

*CLU* is further implicated as an important mediator of TMZ resistance in GBM [95]. M16 is also enriched for targets of acetylcysteine, hymecromone (aka 4-MU), methylthioninium chloride (aka methylene blue) and dequalinium (Figure 4C), all of which have shown promise in pre-clinical studies as GBM therapeutic agents [96–100].

M15 is enriched for mitosis related pathways (Figure 3B). Prior work has identified the hub gene *TYMS* (Figure 3D) as a modulator of TMZ resistance and HMGB2 as a modulator of radioresistance in GBM [101,102]. M15 is enriched for targets (Figure 3C) of the well-established chemotherapeutic agent Gemcitabine, which has been shown to cross the blood-brain barrier and sensitize GBM to radiotherapy [103], in addition to the antioxidant dietary supplement Quercetin, which has shown substantial pre-clinical evidence for treating GBM [104–106].

The most overexpressed modules in MVP include M7 and M17 (Figure 4B). Term enrichment analysis of M7 highlights its distinct role in angiogenesis and ECM organization, consistent with defining features of the MVP region (Supplement Figure 5A). M7 hub genes *TAGLN*, *NOTCH3*, and *ITGA1* (Supplement Figure 5B) have been identified in prior work for their critical involvement in GBM resistance to TMZ [107–109]. The M7 hub gene *FN1* has been identified as an essential gene in *GBP2*-driven GBM invasion [110] and novel therapeutic target [111,112]. M7 is also significantly associated with *COL4A1*-*ITGA1*-*ITGB1* signaling (Supplement Figure 6), supporting a role for this interaction in angiogenesis driven by integrin signaling on endothelial and mural cells in MVP. Further, M7 is enriched for positive targets of the *TBX2* regulon (Figure 4D), which is overexpressed in MVP (Supplement Figure 8) and associated with TMZ resistance [113].

M17 is enriched for response to fibroblast growth factor and negative regulation of focal adhesion assembly (Supplement Figure 5A). Drug target enrichment (Figure 4C) identifies alitretinoin, menadione, and gossypol as potential therapies targeted towards this module.

Retinoic acids have been highlighted in prior work on novel GBM therapies specifically for their role in differentiating GSCs and reducing GBM cell viability [114,115]. Menadione can increase oxidative stress in GBM cells, increasing their vulnerability to redox-disrupting therapies [116,117]. In addition, prior work has evidenced that gossypol reduces growth of TMZ-resistant GBM [118].

The most overexpressed module in PAN is M4 (Figure 4B). M4 is enriched for cellular response to hypoxia and HIF-1 signaling (Supplement Figure 5A). M4 hub genes *HILPDA*, *ADM*, *NDRG1*, *AKAP12*, *SLC2A1*, and *ANGPTL4* (Supplement Figure 5A) are markers for the Neftel MES2 tumor state [50], recapitulating earlier findings of MES2 enrichment in PAN (Supplement Figure 4B). *HILPDA* has also been shown to be central to PAN region specific lipid droplet accumulation, GBM cell state related lipid metabolism modulation, and a potential therapeutic target in GBM [119]. Hub genes *ADM*, *ANGPTL4*, *SERPINE1*, and *CA12* have been identified as actionable targets to overcome TMZ resistance [120–123]. In addition, *AKAP12* has been identified as a biomarker of resistance to *VEGF* targeted therapy [124]. Further, M4 is enriched for targets of arsenic trioxide and celecoxib (Figure 4C). Arsenic trioxide is a chemotherapeutic agent capable of crossing the blood brain barrier with some promise for GBM therapy [125,126]. Celecoxib has evidence across multiple clinical trials indicating its potential benefit as an adjuvant treatment [127]. M4 is enriched for positive targets of *CEBPB*, *CEBPD*, and *FOSL2* regulons (Figure 4D), all of which are overexpressed in PAN (Supplement Figure 8) and have been implicated in driving programs promoting therapy resistance in GBM [127–129].

The most overexpressed modules in LE are M3 and M19 (Figure 4B). M3 is enriched for pathways including neurotransmitter secretion and neuron projection morphogenesis (Supplement Figure 5A), reinforcing prior findings of increased neuronal signaling and synaptic activity in the LE (Supplement Figure 1C). M3 hub genes *STMN2*, *SNAP25*, and *GNG3* (Supplement Figure 5B) associate this module with the Neftel NPC2 tumor state [50], reinforcing prior findings of NPC2 enrichment in LE (Supplement Figure 4B). Ephrin interaction association with M3 (Supplement Figure 6) further supports the role of ephrin signaling in promoting tumor invasion through neuronal guidance, with *EFNB3*-*EPHB6* interacting between NPC-like and neuron cells. Further, M3 is enriched for positively regulated targets of *ARNT2* and negatively regulated targets of *SOX10* regulons (Figure 4D), both of which are overexpressed in LE (Supplement Figure 8). Prior work has shown knockdown of *ARNT2* decreased GSC properties and GBM tumorigenicity programs in vivo [130]. In contrast, suppression of *SOX10*, a consequence of TMZ treatment, promotes GBM progression and a neural stem-cell like cell state [131]. These findings suggest *SOX10* and *ARNT2* may act in opposing directions along the CT-LE gradient, with important implications for GBM behavior and treatment response.

M19 is enriched for neuron differentiation and neuron projection morphogenesis pathways (Supplement Figure 5A). The top M19 hub genes (Supplement Figure 5B) are mitochondrially encoded genes, potentially indicating contamination from cell death. However, recent literature has indicated that in the context of cancer tissue research this may represent meaningful biological signal in relation to metabolic regulatory alterations [132]. In addition, mitochondrially encoded gene expression has been found to be inversely correlated with WHO glioma grade and the M19 hub genes *MT-ATP6*, *MT-CO3*, *MT-CYB*, *MT-CO2*, and *MT-ND4L* have been positively correlated with improved prognosis in glioma [133]. M19 is enriched for targets of metformin (Figure 4C), which has evidence for a potential adjuvant role in a variety anti-cancer therapies including GBM [134–136].

In total, these findings place numerous GBM target genes and therapies in a spatial and functional context, with important implications for the development of rational combination therapies informed by comprehensive niche and pathway coverage.

## Discussion

In this work, we systematically resolved the spatial complexity of GBM by jointly analyzing a large cohort of ST slides using a range of computational pipelines. Our analyses reveal that distinct anatomical niches within GBM impose unique transcriptional and signaling programs on tumor cells, resulting in marked variation across regions. We deconvolved our ST data to extract tumor-specific expression profiles, enabling robust dissection of malignant cell programs despite microenvironmental admixture. The resulting data faithfully recapitulated established spatial trends in GBM biology, including the niche-specific enrichment of canonical gene modules such as those described by Neftel et al [50], with distinct transcriptional states mapping to perivascular, hypoxic, and invasive regions. Comparative analysis of tumor cell expression across these niches revealed extensive gene expression changes, consistent with the well-documented association between spatial microenvironments and GBM cell state heterogeneity [10,12].

We further identified TF modules exhibiting highly localized spatial activity on tumor cell-specific programming. Several TF modules were specifically active in the perivascular niche, each associated with discrete biological processes, while others were restricted to the LE or PAN regions. Notably, we discovered two TF modules with activity concentrated along niche interfaces: one at the CT-PAN boundary, enriched for metabolic adaptation to hypoxia, and another at the CT-LE interface, strongly associated with RNA splicing and isoform diversity, an emerging hallmark of tumor invasiveness in GBM [43]. Importantly, the activity of these 9 TF modules varied across individual slides, reflecting the limited and heterogeneous spatial sampling inherent to each tissue section. This underscores the necessity of pooling data across a large, representative cohort to comprehensively capture the spatial regulatory landscape of GBM [12,15]. Our integrative approach, leveraging a unified data embedding, provides a powerful framework for studying spatially resolved gene regulatory programs and uncovering novel mechanisms of tumor adaptation in GBM.

We further characterized spatial niche-specific cell communication between malignant and non-malignant cell types through a novel integrative workflow combining SC and ST analysis. In the CT, we identify the localization of the *AREG-EGFR* axis mediated by immune cells and targeting AC-like/OPC-like tumor states. Our analysis proposes a novel role for this pair, specifically immune-mediated targeting of AC-like/OPC-like tumor states. We also discovered ECM*-CD44* interactions between mural cells and AC and OPC-like tumor cells. In MVP we identify dual roles of LR signaling, pro-metastatic through angiogenesis driven by integrin signaling, and anti-tumorigenic driven by immune signaling via *GZMA-F2R*. In PAN, we uncovered spatially patterned mechanisms of the inflammatory response, specifically immunomodulation driven by *MIF*, and MES-like state transition driven by *FN1/SPP1-CD44* interactions. *MIF* signaling is proposed to have a complex role in GBM, and our analysis supports the role of *MIF* in driving vascular mimicry [77,137]. In LE, we identify ephrin signaling driven neuronal guidance and tumor invasion through interactions between NPC-like tumor cells and neurons.

In addition, we comprehensively characterized molecular pathway patterning across spatial niches through the lens of gene co-expression network modules. These modules reinforce and are reinforced by established spatial patterns of biological pathway activity in GBM and highlight region specific associations for Neftel tumor states. Further, they place independently studied GBM therapeutic targets and mediators of therapy resistance into spatial functional contexts based on their hub gene status or regulatory relationship with modules and the module’s association with Ivy GAP niches. In addition, for each region we are able to identify pathways enriched for a drug or drug class with promising evidence in the GBM literature. Together, these findings help to inform rational combination therapies based upon actionable targets across the diverse pathway and histological niche landscape of GBM.

Despite these advances, some limitations remain. The resolution of Visium spatial profiling constrains our ability to pinpoint tumor-specific expression changes at SC resolution, and our deconvolution depends on the breadth and quality of available SC references. We also have not fully characterized the dynamic states of immune and stromal cells across niches.

Recent studies have highlighted extensive immune rewiring in the GBM ecosystem [138] and the important functions of cancer-associated fibroblasts in tumor growth and treatment response [139]. Future work employing higher-resolution spatial profiling will be crucial for unraveling the full complexity of the tumor microenvironment and understanding how spatial context shapes cell states. In addition, further work designed to explore the temporal nature of these niche characteristics, including their response to treatment and transformations during recurrence, will shed further light on our findings’ clinical implications.

Our findings have direct therapeutic implications. The remarkable diversity of cell states within a single tumor likely explains the limited efficacy of single-agent therapies, such as *EGFR* or *VEGF* inhibitors. This work underscores the urgent need for rational combination therapies that can simultaneously target key vulnerabilities or master regulators within each tumor niche.

Embracing the spatial and functional heterogeneity of GBM will be essential for developing more effective and durable treatment strategies for this devastating disease.

## Methods

### 1. Sample Collection

Under an institutional review board approved study in accordance with recognized ethical guidelines, intraoperative brain tumor specimens were collected at the University of Michigan using MRI-imaged guidance technology to localize specimen by senior author WNA. These specimens were obtained after written informed consent from patients. Fourteen tissue sections were collected in house from 7 patients. Five slides included adjacent sections of the same tumor tissue, and slides 1512-A/B and 1512-C/D were collected from different regions of the same tumor. Eight slides are primary tumor, while six (5492-C/D, 8379-A/B/C/D) were from recurrent GBM. Freiburg cohort Visium ST data were downloaded from DRYAD (https://datadryad.org/dataset/doi:10.5061/dryad.h70rxwdmj) and the Broad cohort Visium ST data was downloaded from zenodo (https://zenodo.org/records/12624860).

### 2. Processing and Quality Control of Visium ST Data

Spots with fewer than 200 expressed genes were labeled as low quality and excluded from downstream analyses. Similarly, spots with over 5% hemoglobin genes were excluded due to excessive bleeding in tissue. Any slide with fewer than 100 spots passing quality-control (QC) thresholds was not used for integrative analysis. Slides passing QC for each cohort were independently merged into a single *Seurat* [140] object. Count data analysis was carried out using *Seurat v5.2* [140]. Raw counts were normalized using the *NormalizeData* function. The top 3000 variable genes were used for dimension reduction using PCA with the *RunPCA* function, and the top 15 PCs were used to construct a UMAP data embedding using *RunUMAP* for sample visualization in the UM cohort.

### 3. Spot-level Spatial Niche Scoring for Gene Sets

Normalized counts per spot were used to compute enrichment scores for different gene sets of interest using *UCell* [16]. This algorithm was specifically developed for dealing with data sparsity in single cell datasets and uses the non-parametric Mann-Whitney U statistic for robust computation of enrichment scores. Scores are computed using the *AddModuleScore_UCell* function with maxRank of 3000. Spot-level niche annotation was carried out by computing enrichment scores for regional markers from the Ivy GAP study [7], and labeling spots based on maximal scores. The same algorithm is used for computing drug target pathway and cancer hallmark pathway scores per spot, with gene sets downloaded using *msigdbr* [141]. Drug target pathways by gsea-msigdb standard name were PID_AVB3_INTEGRIN_PATHWAY, KEGG_MEDICUS_REFERENCE_AREG_EGFR_PI3K_SIGNALING_PATHWAY, PID_VEGF_VEGFR_PATHWAY, and PID_PDGFRA_PATHWAY.

### 4. Differential Expression Analysis

Differential expression analysis with genes and pathway scores was carried out using the *FindAllMarkers* function in *Seurat*, with the default Wilcoxon Rank-Sum test for the appropriate assay object between conditions of interest. Unless otherwise stated, we used significance thresholds of FDR < 0.05 and log2 fold-change > 1 to identify significant markers. Enrichment analysis for significantly differential genes was carried out using the *compareCluster* function form *clusterProfiler* [142,143].

### 5. Visium Spot Deconvolution

Visium ST spot deconvolution was performed with *BayesPrism* [34] using a comprehensive GBM single-cell atlas [47] as reference. We used the level-3 annotation from the atlas for cell type cell state annotation. We downsampled the atlas to 3000 randomly selected cells per reference group. Raw counts for protein-coding genes from single-cell and Visium ST data were used to create the *BayesPrism* object. For the inference, we used *InstaPrism* [48], which re-implements *BayesPrism* by replacing the Gibbs sampling step with a fixed-point algorithm, providing considerable speed and memory advantages. In addition to ST spot compositions, we also use the algorithm to extract spot-level tumor-specific expression data for the whole transcriptome.

### 6. Constructing Cell-Interaction Graph

The inferred spot-level cell type compositions from deconvolution are used to construct a cell-cell interaction graph. Since the data is compositional, we use the centered-log ratio transform to bring them into Euclidean space, and then use the graphical-lasso algorithm [144] to infer an interaction graph between cell types. We show all non-zero edges at a sparsity threshold of lambda 1, since the suggested lambda_min_ and lambda_lse_ values gave too many interactions.

### 7. Learning 2D PCA Embedding to Capture Intrinsic Spatial Context Across Slides

To study how biological processes are continuously changing along spatial gradients, we learned a 2D data embedding that preserves the spatial tissue context of Visium ST spots across slides. We trained an *scVI* model [145] across all 44 slides for the whole transcriptome, treating each slide as an independent batch. We used a model with two layers and 30 latent variables, with the negative binomial likelihood for modeling gene expression counts. The trained model was used to generate counterfactual normalized, batch-corrected counts (mapped to reference slide 1512-A) for the Ivy GAP marker genes for all spots across the entire dataset. These batch-corrected counts were used to generate a supervised PCA-embedding for the dataset.

### 8. Hotspot Analysis to Identify TF Modules

We used *Hotspot* [37] to identify modules of spatially co-expressed TFs from the PC embedding. We construct a kNN graph on the PC embedding with k=15 to compute auto and cross-correlation metrics; and use the tumor-specific expression data to identify modules of TFs with spatially localized activity in tumor cells. Autocorrelation test identified 184 out of 1457 tested TFs as showing significant spatially varying patterns of expression. We then compute cross-correlation metrics between these TFs and run hierarchical clustering to identify modules with at-least 10 genes in each. Spot-level activity of each TF module is computed using the *calculate_module_scores* function and used for visualization in spatial coordinates and PC embedded space.

We identified the top 1000 spots with the highest score for each TF module, and compared the tumor-specific gene expression program in these spots as described earlier, followed by gene ontology [17] enrichment analysis. For module 9, we specifically contrasted its expression profile against the top scoring cellular tumor and leading edge spots to identify unique molecular programs specifically enriched in this domain.

### 9. High Dimensional Weighted Gene Co-expression Network Analysis

High dimensional weighted gene co-expression network (GCN) analysis was performed on the Visium ST data using *hdWGCNA v0.4.05* [88]. To address the sparsity inherent to Visium ST data, counts from spots passing QC were aggregated by spatial proximity into metaspots through summation. Metaspots only included spots from the same slide and IVY region. A minimum expression threshold was applied to metaspots, retaining genes with count greater than 1 in at least 20% of samples. Metaspot counts were normalized and log transformed. Outlying metaspots were removed using the standardized connectivity-based method implemented in *BioNERO v1.12.0* [146] with the standardized connectivity threshold, zk, set to -3.

The resultant filtered and log-normalized expression data was passed on to network construction. All network construction steps were performed using the biweight midcorrelation function and signed hybrid network type where applicable. The scale-free topology model fit for soft power thresholding values ranging from 4 to 16 was assessed and the lowest soft power achieving a fit of 0.80 was used for network construction. GCN construction and module detection was performed using the *ConstructNetwork* function in *hdWGCNA* with a minimum module size cutoff of 20 genes. Modules with module eigengene (MEs) dissimilarity less than 0.15 were merged. ME calculation included correction for the slide of origin using the Harmony algorithm [147].

Hub genes were identified by their correlation with their ME (kME), where the larger gene set of either the top 10 or 10% of module genes were selected by kME. Module enrichment for pathways, ontology terms, and drug targets was performed using *enrichR v3.4* [148,149]. To focus on core module functionality, only up to the top 300 genes as sorted by kME were included in enrichment analysis for each module. The association between module expression levels and IVY region annotation was performed in two ways. First, for each region, one-vs-all testing was performed on MEs using the *FindAllDMEs* function in *hdWGCNA*, which performs Wilcoxon rank-sum testing between MEs from spots in one region against the MEs from spots in all other regions. Second, a linear mixed effect model was implemented using *lmerTest v3.1-3* [150]. For each module, the ME was modeled against the one-hot-encoded IVY region assignment with a random intercept included for the slide.

Transcription factor regulatory network (TRN) analysis was performed following the approach in Childs & Morabito et al. [151] using *hdWGCNA v0.4.05* [88]. Candidate TF target genes were identified in the gene co-expression network by the presence of TF binding motifs in gene promoter regions using the JASPAR database [152] of TF binding profiles and *motifmatchr v1.26.0* [153]. TF regulons were defined as up to the top 50 target genes for each TF with a regulatory score of at least 0.05. For each regulon, enrichment analysis of positively and negatively regulated TF targets in gene co-expression network modules was performed using the *fgsea* function from *fGSEA,* with gene scores corresponding to TF-target association in the regulon.

### 10. Cell Communication Inference

To identify ligand-receptor (LR) pairs playing an active role in each Ivy GAP region, we developed a novel workflow integrating both SC and ST data. We start by performing two analyses in parallel: identifying top interacting LR pairs using *CellChat* [52] and the GBMap single cell atlas [154] with level 3 annotations, and identifying LR pairs whose spatial co-expression is significantly associated with the Ivy GAP region. To perform the first SC analysis, *CellChat* uses a mass action model to identify significantly interacting LR pairs between all possible cell type combinations. We performed global p-value adjustment using Benjamini-Hochberg (BH) to correct for false positives.

To perform the second spatial analysis, we first calculated an LR co-expression metric over all 44 samples for those samples that had more than 1% of spots with non-zero ligand and receptor expression. For contact-based LR signaling, we simply calculated the product of ligand and receptor expression within the spot. For diffusion-based LR signaling, we calculated the spot-level bivariate Moran’s statistic with a bandwidth of 1 radius away from the center spot. We then fit a mixed-effect model to represent the spatial co-expression scores as a function of the Ivy GAP region score using the sample as a random effect. We performed p-value adjustment through BH across all LR pairs for each region, selecting the pairs with significant and positive coefficient for that region. These pairs are interpreted to have increased co-expression deeper within the region of interest, that is, a likely higher interaction within this region.

We used *commot* [155] to create all spatial co-expression maps for validation. Commot uses optimal transport with the consideration that multiple ligands can bind to one receptor, and vice versa. While computationally inefficient for evaluating a large number of LR pairs, this method can identify both signaling strength and direction, while basic co-expression statistics (such as cosine similarity or bivariate moran’s) cannot. This allows us to visualize strong localized signaling to validate region-specific cell-cell interactions found by our quantitative workflow.

### 11. Association of LR Pairs with Gene Co-expression Network Modules and Functional Pathways

To infer functional roles of top LR pairs per region, we integrated cell-cell interaction with gene co-expression network analysis, modeling LR spatial co-expression as a function of module eigengene scores with sample as random effect. We performed p-value adjustment using BH across all modules for each LR pair, and ran this analysis for pairs unique to CT and MVP, as well as all pairs spatially associated with LE and PAN. Through a literature review on the LR pairs associated with modules specific to the region, we can then hypothesize LR drivers of module mechanisms.

### 12. Data and Code Availability

The SC and ST data generated in this study are provided as Python AnnData objects through the following link: https://zenodo.org/records/16505469. Access to all code used in this analysis is available upon reasonable request.

## Acknowledgements

We would first and foremost like to thank all of the tissue donors who made this work possible. The work was supported by NINDS (https://www.ninds.nih.gov/) K08 NS128271, NCI (https://www.cancer.gov/) P50-CA-269022, University of Michigan Medicine Chad Carr Pediatric Brain Tumor Center Research Award and grants from the B*Cured Foundation (https://bcured.org/). Additionally, RK was funded by the University of Michigan Biomedical Informatics and Data Science Training Program (T32-GM141746). In addition, AM received funding through the NIH grants F071759 and UM1DK126185. TH was supported by University of Michigan Rogel Cancer Center, Brain Research Foundation, American Brain Tumor Association, Seth M Boyer Foundation, V Foundation for Cancer Research, and Department of Defense Glioblastoma Research Program HT94252510960. The funders had no role in study design, data collection and analysis, decision to publish, or preparation of the manuscript. The authors do not have any competing interests or conflicts of interest related to this presented work.

## Supplementary Tables and Figures

**Supplement Figure 1.**
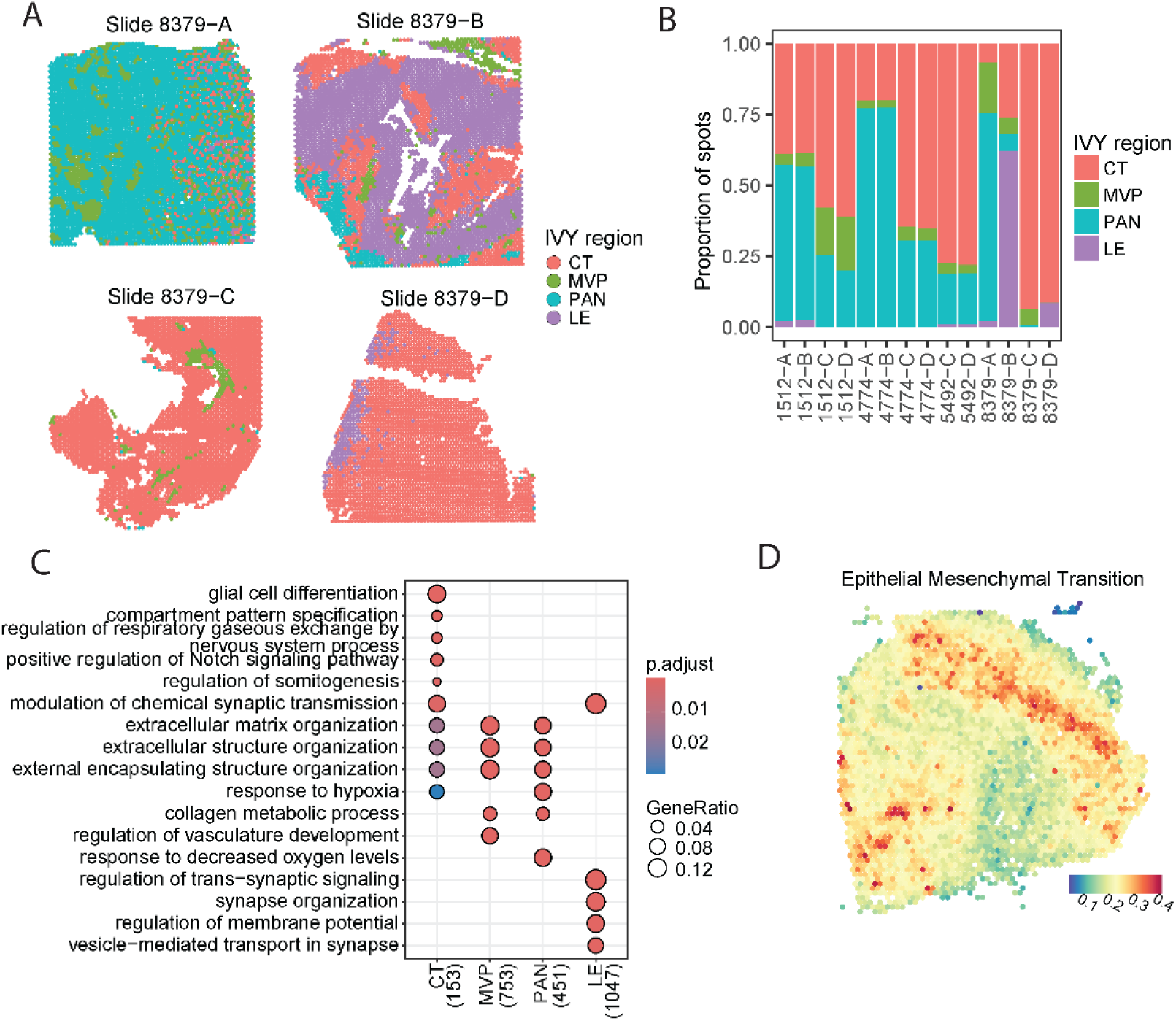
: Supplement to Figure 1. **(A)** Representative niche annotations for additional ST slides in our cohort. **(B)** Relative niche compositions of the 14 ST slides in our cohort. **(C)** GO term enrichment for differentially expressed genes across niches. **(D)** Spatial localization of EMT signature to MVP niche in representative slide.

**Supplement Figure 2.**
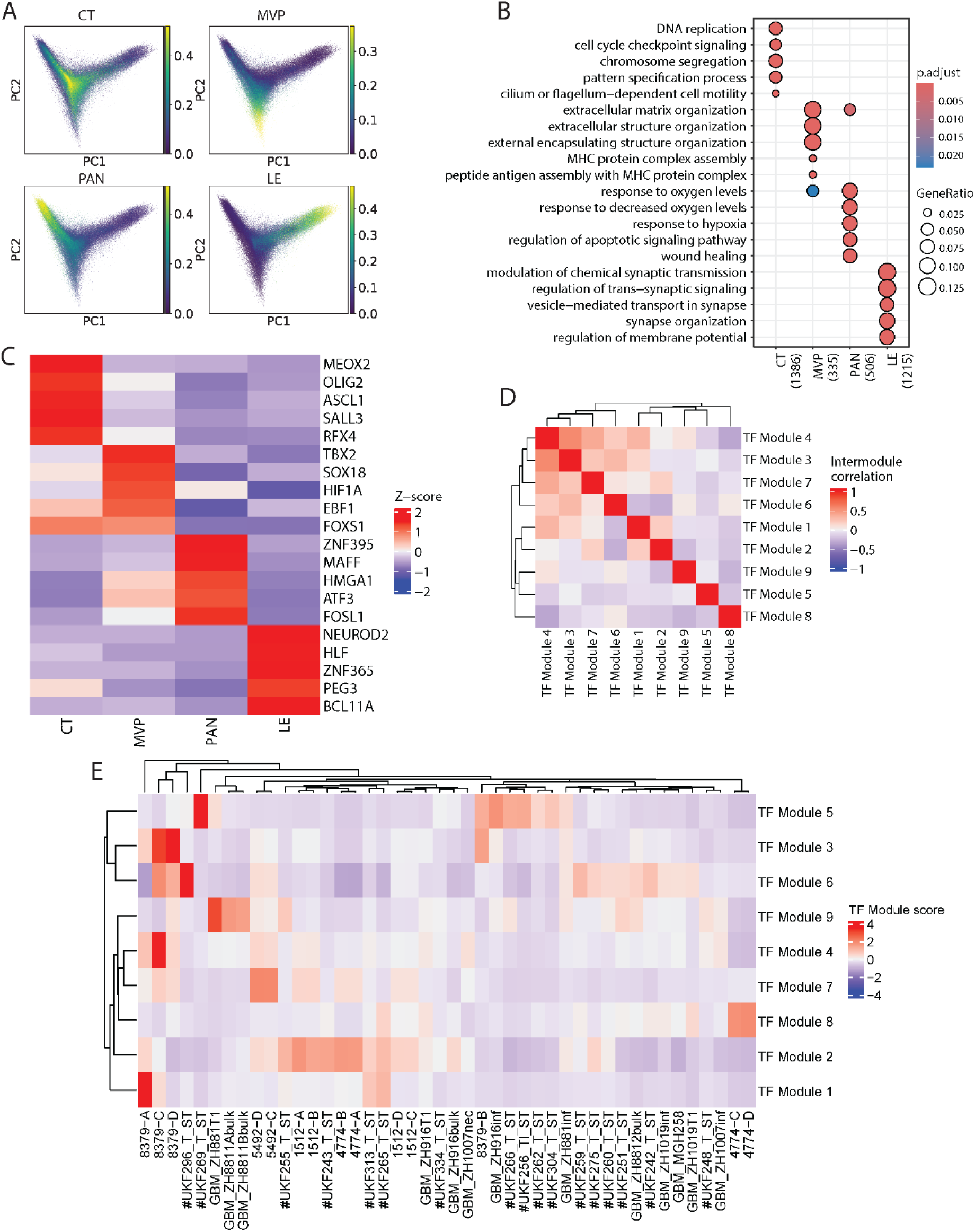
: Supplement to Figure 2. **(A)** 2D PCA embedding captures continuous spatial gradients across multiple slides**. (B)** GO Enrichment analysis of tumor-specific differential expression test between GBM niches. **(C)** Scaled expression of Top-5 differentially expressed transcription factors in each niche. **(D)** Correlation plot between different TF modules shows spatial segregation of their activity**. (E)** Mean activity of TF modules by slide shows importance of using large cohort to identify distinct spatially localized programs in the disease due to inter-patient variance in spatial niche sampling.

**Supplement Figure 3.**
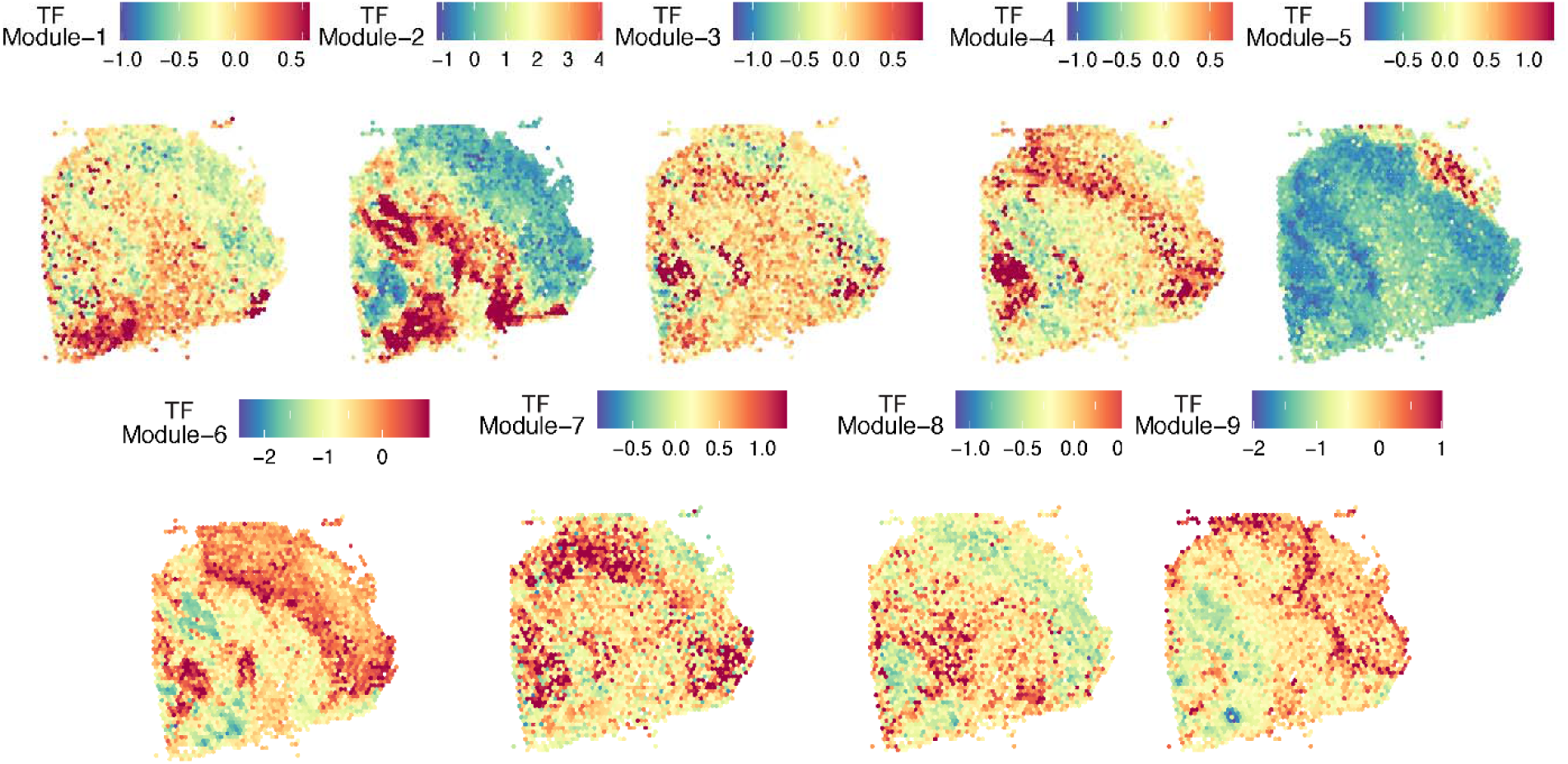
: Spatial activity of TF modules identified with Hotspot analysis in representative slide 1512-A.

**Supplement Figure 4.**
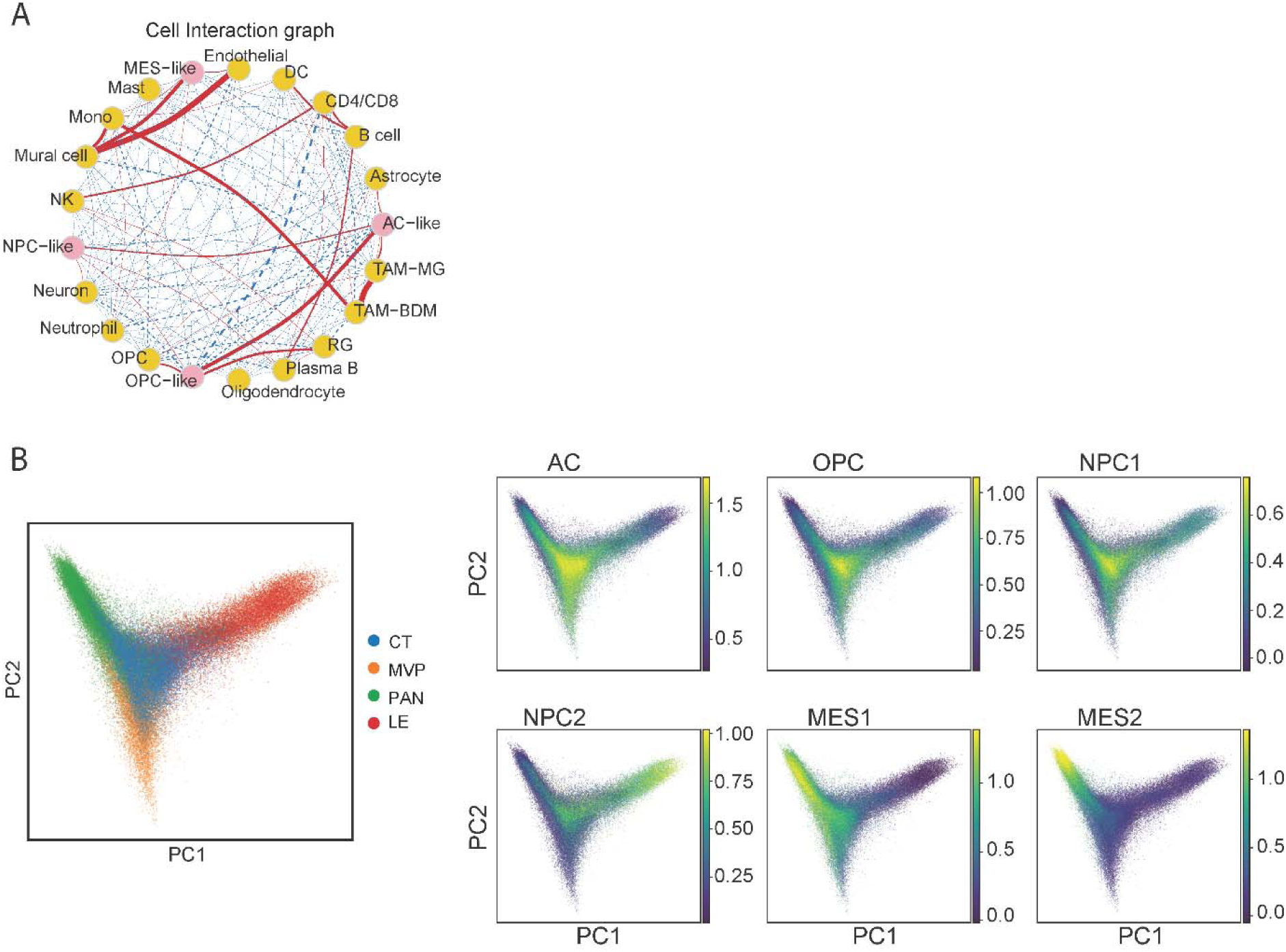
: Supplement to Figure 3. **(A)** Cell-interaction graph. Red lines indicate co-occurrence; blue dotted lines indicate repulsion. **(B)** Localization of Neftel tumor state enrichment to specific domains in embedding space. 2D PCA embedding captures continuous spatial gradients across multiple slides.

**Supplement Figure 5.**
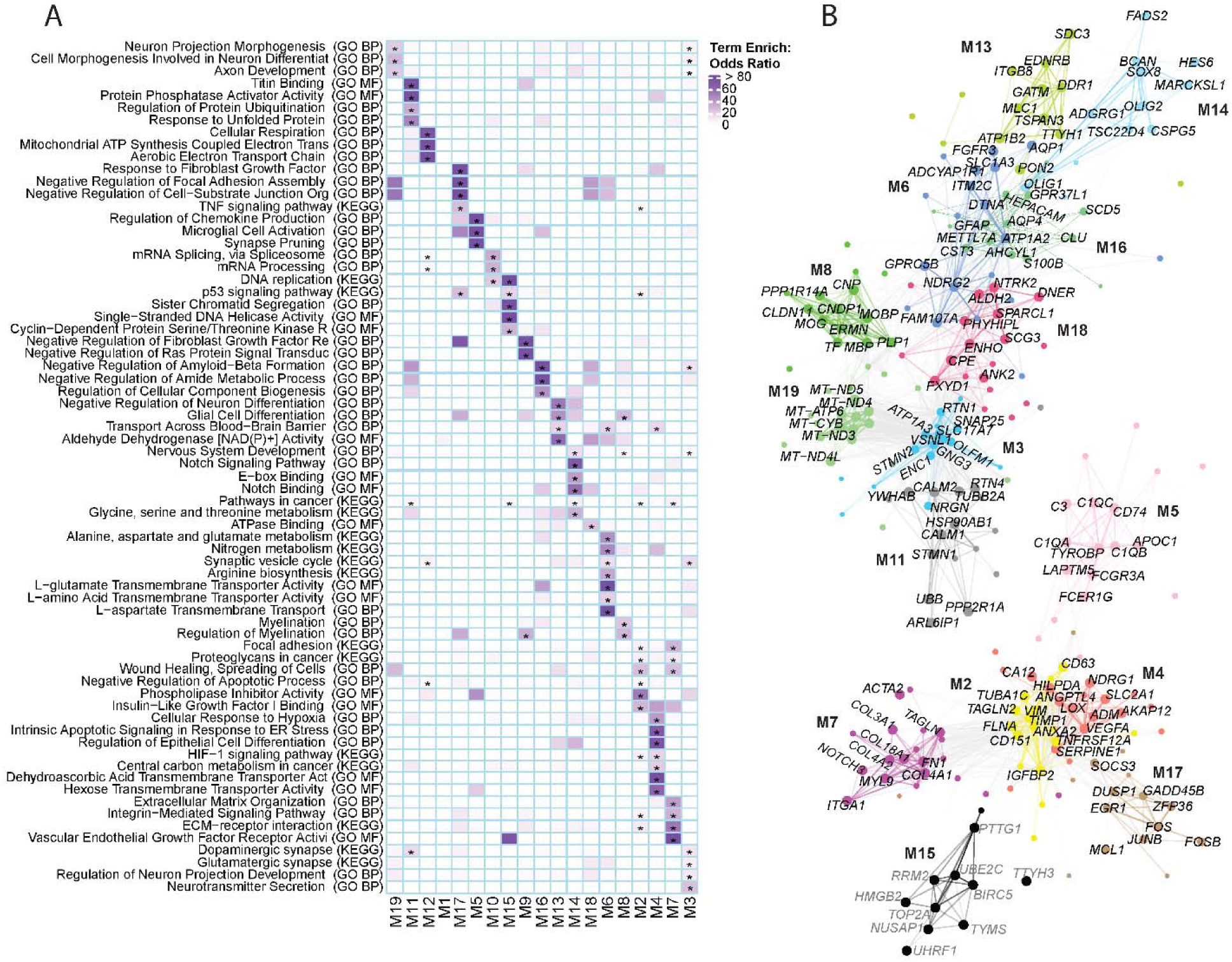
: Supplement to Figure 4. **(A)** Selection of leading significant (FDR < 0.05) pathway enrichment results in each module. **(B)** Visualization of network, subset to select niche-associated modules. Top-10 module hub genes labeled for each module.

**Supplement Figure 6.**
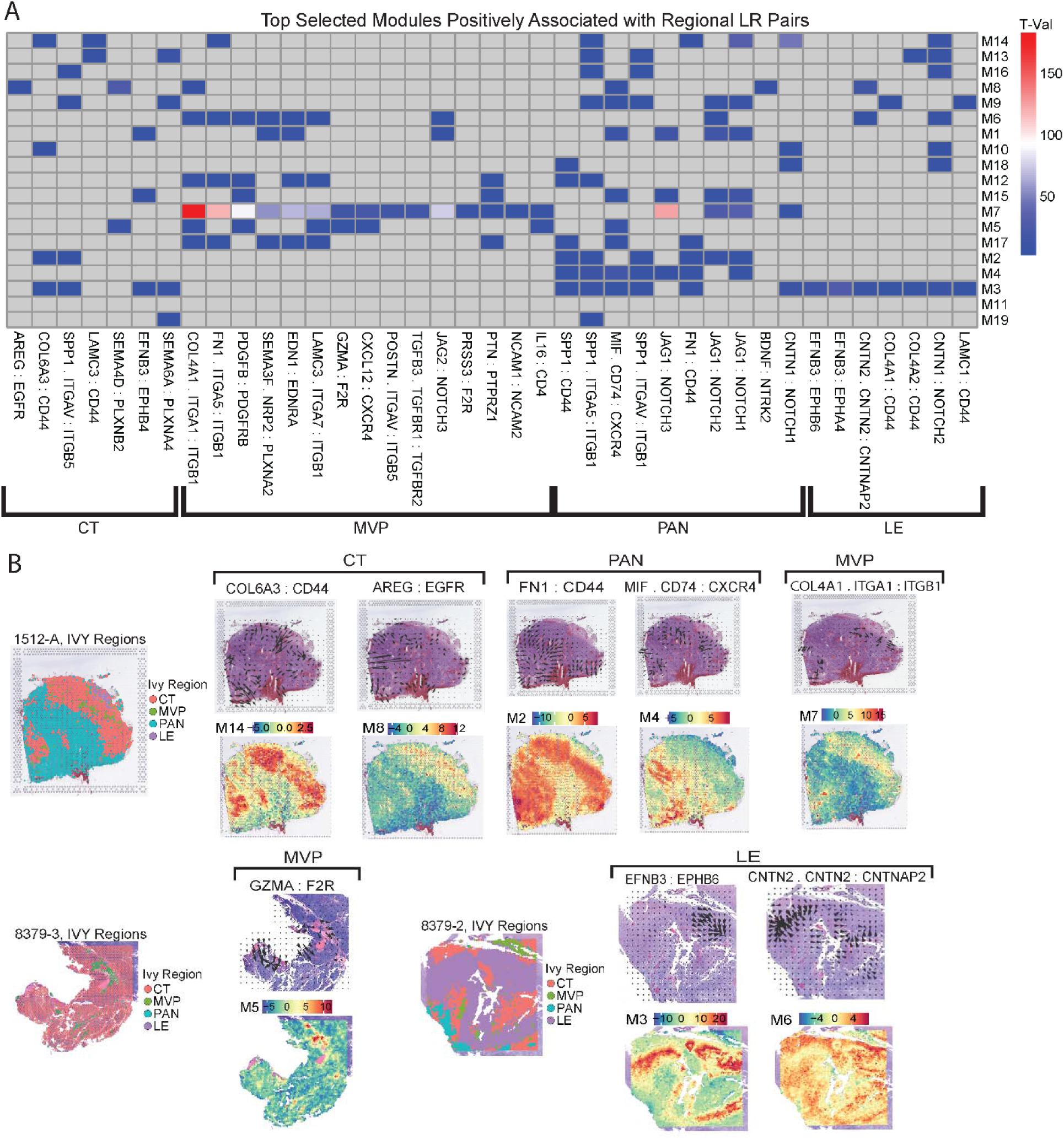
: Ligand-receptor analysis reveals distinct signaling and interaction-based drivers of spatial module behavior. **(A)** T-values from regression of ligand-receptor co-expression against module eigengene. Showing selection of most significant results for each IVY Gap region. Insignificant relationships in gray. **(B)** Visualization of commot sender statistics for select ligand-receptor pairs and module eigengenes in representative slides for each IVY Gap region.

**Supplement Figure 7.**
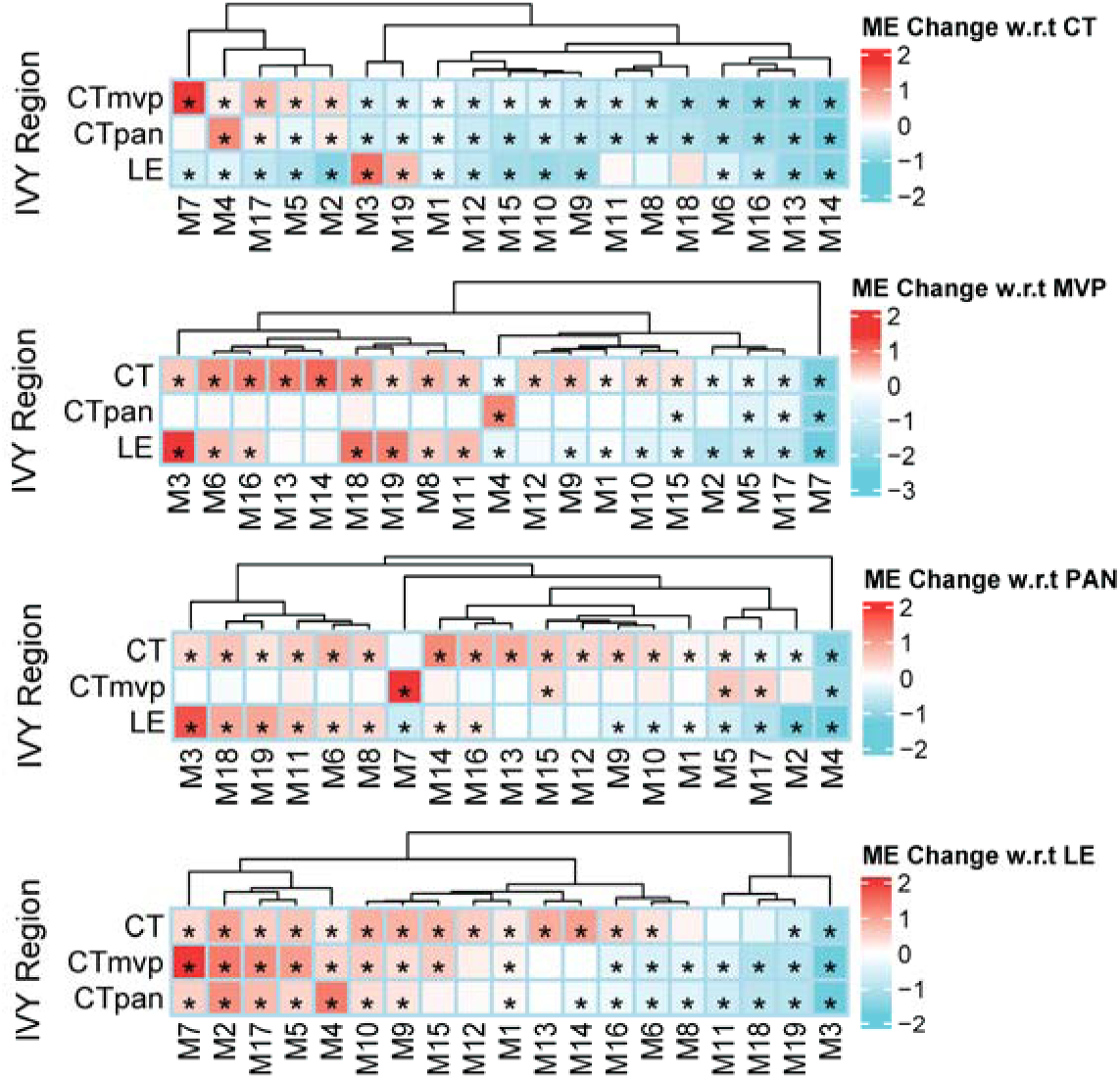
: Linear mixed effect modeling of module eigengenes in relation to IVY Gap niche. Heatmap shown for each IVY Gap niche (CT, MVP PAN, LE) as reference. * where FDR < 0.05.

**Supplement Figure 8.**
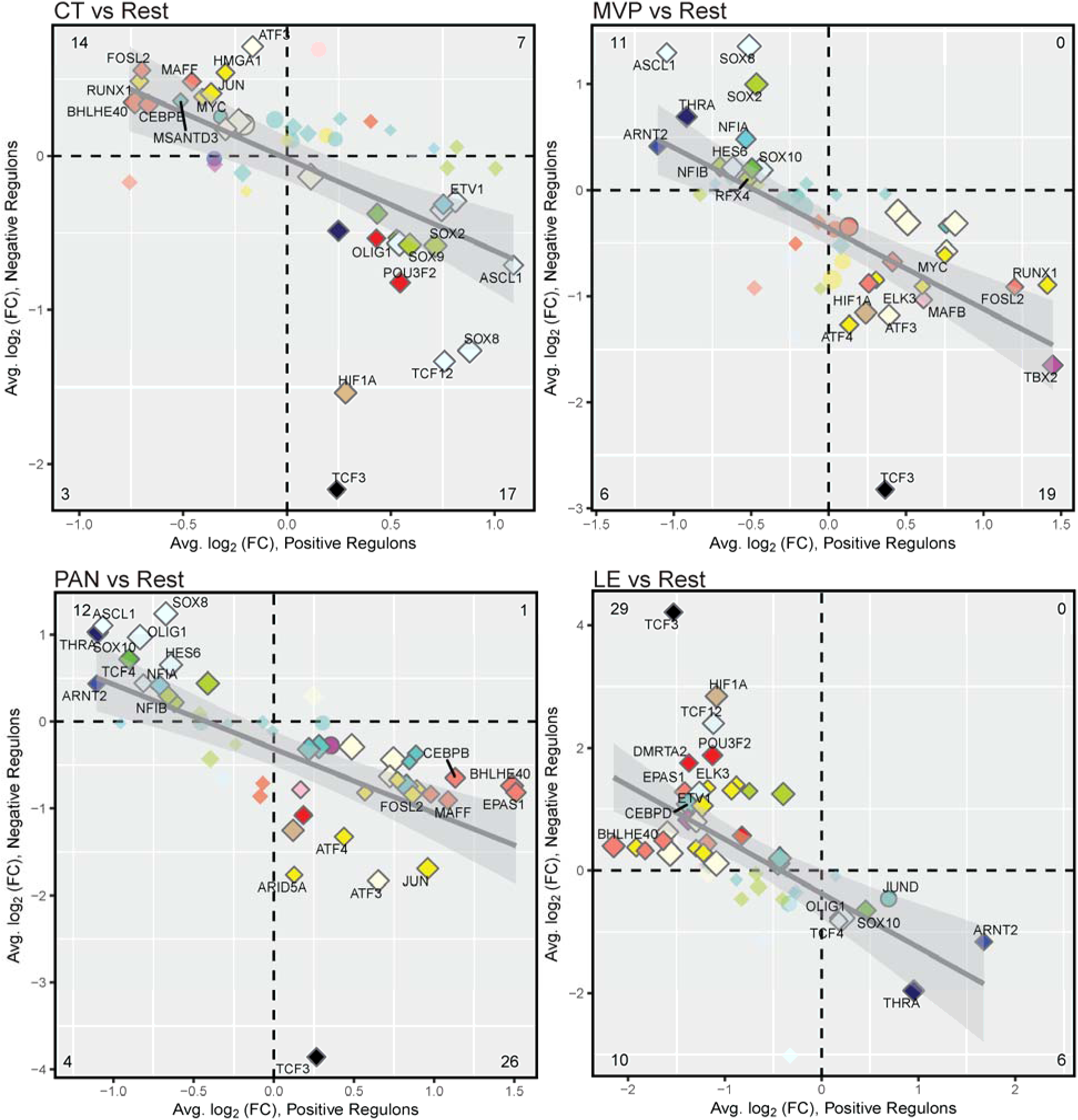
: *Differential regulon analysis*. Analysis shown for each IVY Gap niche (CT, MVP, PAN, LE). Each point represents a TF colored by module assignment in the co-expression network. Diamonds represent TFs that are also significantly differentially expressed with respect to niche. Regulons that did not reach significance are opaque, while significant regulons have a gray outline. The number of significant differentially expressed regulons in each quadrant is labeled in figure corners.

